# Fully Human Monoclonal Antibodies Effectively Neutralizing Botulinum Neurotoxin Serotype B

**DOI:** 10.1101/2019.12.18.881920

**Authors:** Takuhiro Matsumura, Sho Amatsu, Ryo Misaki, Masahiro Yutani, Anariwa Du, Tomoko Kohda, Kazuhito Fujiyama, Kazuyoshi Ikuta, Yukako Fujinaga

## Abstract

Botulinum neurotoxin (BoNT) is the most potent natural toxin known. Of the seven BoNT serotypes (A to G), types A, B, E, and F cause human botulism. Treatment of human botulism requires the development of effective toxin-neutralizing antibodies without side effects such as serum sickness and anaphylaxis. In this study, we generated fully human monoclonal antibodies (HuMAbs) against serotype B BoNT (BoNT/B1) using a murine–human chimera fusion partner cell line named SPYMEG. Of these HuMAbs, M2, which specifically binds to the light chain of BoNT/B1, showed neutralization activity in a mouse bioassay (approximately ≥ 100 i.p. LD_50_/mg of antibody), and M4, which binds to the C-terminal of heavy chain, showed partial protection. The combination of M2 and M4 was able to completely neutralize BoNT/B1 with a potency greater than 10,000 i.p. LD_50_/mg of antibodies, and was effective both prophylactically and therapeutically in the mouse model of botulism. Moreover, this combination showed broad neutralization activity against three type B subtypes, namely BoNT/B1, BoNT/B2, and BoNT/B6. These data demonstrate that the combination of M2 and M4 is promising in terms of a foundation for new human therapeutics for BoNT/B intoxication.

**IMPORTANCE:** Botulinum neurotoxins (BoNTs) produced by *Clostridium botulinum* and related species cause human botulism. Immunotherapy is the most effective treatment for botulism and equine immune serum formulations are used in cases of human botulism. However, these antisera may cause serum sickness or anaphylaxis. Additionally, the production of immune sera involves complicated and time-consuming manufacturing processes and quality management. Therefore, the development of safe, effective, and higher productive antibodies is required. Here we generated fully human monoclonal antibodies against serotype B BoNT (BoNT/B). We found that the combination of these antibodies (M2+M4) had potent and broad neutralization activity against BoNT/B, and showed therapeutic and preventive effects against botulism in mouse models. These data indicate that M2+M4 are promising candidates for the development of human therapeutics and prophylactics for BoNT/B intoxication.

## INTRODUCTION

Botulinum neurotoxins (BoNTs) produced by the anaerobic bacterium *Clostridium botulinum* and related species cause botulism, a neuroparalytic disease with high mortality (1,2). BoNTs have been classified as category A agents by the Centers of Disease Control and Prevention (CDC) and are listed among the six agents at highest risk of being used as bioweapons (3). Seven serotypes, designated A to G, have been identified, and four of these, namely A, B, E, and F, cause human botulism^2^. It was recently reported that BoNT/H is produced by the novel *C. botulinum* strain IBCA10-7060, which also produces BoNT/B (4,5), and subsequent studies have suggested that BoNT/H is a hybrid-toxin of BoNT/A1 and BoNT/F5 (6–8).

Each BoNT is synthesized as a single polypeptide chain (150 kDa) that is proteolytically activated by cleavage into a light chain (L chain, 50 kDa) and a heavy chain (H chain, 100 kDa), which are linked by a disulfide bond^1^. The L chain acts as a zinc metalloprotease. The H chain consists of two functionally distinct regions: the C-terminal, or receptor-binding, domain (H_C_), and the N-terminal, or translocation, domain (H_N_).

Food-borne and infant botulism are the primary forms of human botulism (9), and are caused by intestinal absorption of BoNT. BoNT in the gastrointestinal lumen crosses the intestinal barrier (10), enters the blood stream, and reaches the neuromuscular junction. There, BoNT binds via HC to the receptors present on presynaptic nerve terminals. BoNT/A, BoNT/D, BoNT/E, and BoNT/F bind to synaptic vesicle protein 2 (SV2) and polysialogangliosides (11–13), whereas BoNT/B and BoNT/G bind to synaptotagmin and polysialogangliosides (14,15); all serotypes subsequently enter neuronal cells by endocytosis. In the acidified synaptic vesicles, HN induces translocation of the L chain into the cytosol (16,17). The metalloprotease domain of the L chain cleaves the soluble *N*-ethylmaleimide–sensitive fusion protein attachment protein receptors (SNAREs) required for synaptic vesicle fusion. The synaptosomal associated protein 25-kDa (SNAP-25) is the target of BoNT/A, BoNT/C, and BoNT/E; synaptobrevin (also known as vesicle-associated membrane protein, VAMP) is the target of BoNT/B, BoNT/D, BoNT/F, and BoNT/G; and syntaxin is the target of BoNT/C (1,2). Cleavage of SNARE inhibits release of the neurotransmitter acetylcholine and leads to paralysis (1,2).

Immunotherapy is the most effective treatment for BoNT intoxication. Equine immune serum formulations are used in cases of human botulism: Botulismus-Antitoxin Behring (Novartis Vaccines and Diagnostics GmbH and Co. KG) is used for treatment of types A, B, and E, while Botulinum Antitoxin Heptavalent (BAT, Cangene Corporation) is used for treatment of types A, B, C, D, E, F, and G. However, because both drugs use heterologous equine proteins, these antisera may cause serum sickness or anaphylaxis. A human immune serum formulation, named BabyBIG (California Department of Public Health) recently became available, but in limited supplies and to treat only infant botulism (9,18). Additionally, the production of immune sera involves complicated and time-consuming manufacturing processes and quality management. Therefore, the development of safe, effective, and higher productive antibodies is required.

In this study, we generated fully human monoclonal antibodies (HuMAbs) against BoNT/B1, designated M2, M4, and S1, using a murine–human fusion partner cell line named SPYMEG (19–21). We found that M2 and S1 bound to the L chain of BoNT/B1, and M4 bound to the H_C_. Furthermore, M2 showed potent neutralization activity, and the combination of M2 and M4 showed a synergistic neutralization effect. Moreover, this combination provided complete protection against BoNT/B1 in models of both botulism treatment (post-exposure) and prevention (pre-exposure). Finally, we confirmed that this combination also possessed neutralization activity against subtypes BoNT/B2 and BoNT/B6. These results indicate that the combination of M2 and M4 may serve as an effective therapeutic agent for BoNT/B intoxication.

## RESULTS

### Immunization with tetravalent botulinum toxoid vaccine

Two healthy adult volunteers were inoculated 4 or 5 times with a tetravalent botulinum toxoid. At 9 and 18 days after the last vaccination, peripheral blood samples were collected from each volunteer. Plasma antibody titers against BoNT/A1 or BoNT/B1 were measured by enzyme-linked immunosorbent assay (ELISA). In both volunteers, the antibody titers were higher than that of non-vaccinated volunteers as previously reported (22) (Table 1).

**Table 1.** Antibody titers against BoNT/A1 or BoNT/B1 in plasma samples from volunteers. Antibody titers against BoNT/A1 or BoNT/B1 from the plasma samples of human volunteers were tested using ELISA. Plasma samples were serially diluted two-fold and 50 μl of each dilution was added to plates coated with BoNT/A1 or BoNT/B1. After washing, bound HuMAbs were detected by anti–human IgG antibody conjugated with HRP. ELISA titers are expressed as the highest dilution factor with an absorbance at least twice that of negative control plasma.

| Volunteer<br>number | Number of<br>vaccinations | Days after last<br>immunization | ELISA titer<br>(log<sub>2</sub>)<br>BoNT/A1 | ELISA titer<br>(log<sub>2</sub>)<br>BoNT/B1 |
| --- | --- | --- | --- | --- |
| Bkf002 | 5 | 9 | 14 | 13 |
| Bkf002 | 5 | 18 | 13 | 15 |
| Bkf003 | 4 | 9 | 13 | 12 |
| Bkf003 | 4 | 18 | 13 | 14 |

### Preparation of HuMAbs

PBMCs were isolated from peripheral blood samples and fused with SPYMEG cells. After HAT selection, ELISA (first screening) was performed using plates coated with a mixture of BoNT/A1 and BoNT/B1. Twenty-seven and eight positive wells, respectively, were obtained from blood samples collected 9 and 18 days after the last vaccination. These wells were subjected to cell cloning by limiting dilution. Finally, we obtained eight stable hybridoma clones. Isotype analysis showed that four hybridoma clones, designated M1, M2, M4, and S1, were produced IgG; three hybridoma clones, designated M3, M5, and M6, were produced IgM; and one hybridoma clone, designated M7, was produced IgA. Furthermore, an IgG subclass assay revealed that M2, M4, and S1 were each comprised of an IgG1 H chain and L (lambda) chain (Table 2).

**Table 2.** Isotypes of HuMAbs *The IgG subclass of M1 could not be determined with the IgG subclass human ELISA kit.

| Volunteer number | Days after last immunization | Clone No, | Isotype |
| --- | --- | --- | --- |
| Bkf002 | 9 | M1 | IgG* |
|  | 9 | M2 | IgG1 |
|  | 9 | M3 | IgM |
|  | 9 | M4 | IgG1 |
|  | 9 | M5 | IgM |
|  | 9 | M6 | IgM |
|  | 18 | M7 | IgA |
| Bkf003 | 9 | S1 | IgG1 |

### Binding specificity of HuMAbs

We selected three IgG1 HuMAbs, namely M2, M4, and S1, and analyzed the binding activity of these HuMAbs against BoNT/A1 and BoNT/B1 by ELISA. All three HuMAbs bound to BoNT/B1, whereas none bound to BoNT/A1 (Fig. 1). We next performed ELISA using recombinant proteins to determine whether the HuMAbs recognized the L and H chains (HN, HC) of BoNT/B1. M2 and S1 bound to the L chain of BoNT/B1, while M4 bound to the HC. All three HuMAbs did not bind to HN (Fig. 2A). To further investigate if the epitopes recognized by HuMAbs overlapped, we performed competitive ELISA. Non-labeled M2 and S1 were equally effective at inhibiting the binding of HRP-labeled M2 to BoNT/B1 and that of binding of HRP-labeled S1 to BoNT/B1, suggesting that M2 and S1 recognized overlapping epitopes. By contrast, binding of HRP-labeled M4 was not inhibited by non-labeled M2 and S1, indicating that M4 recognized a different epitope than M2 and S1 (Fig. 2B).

**Figure 1.**
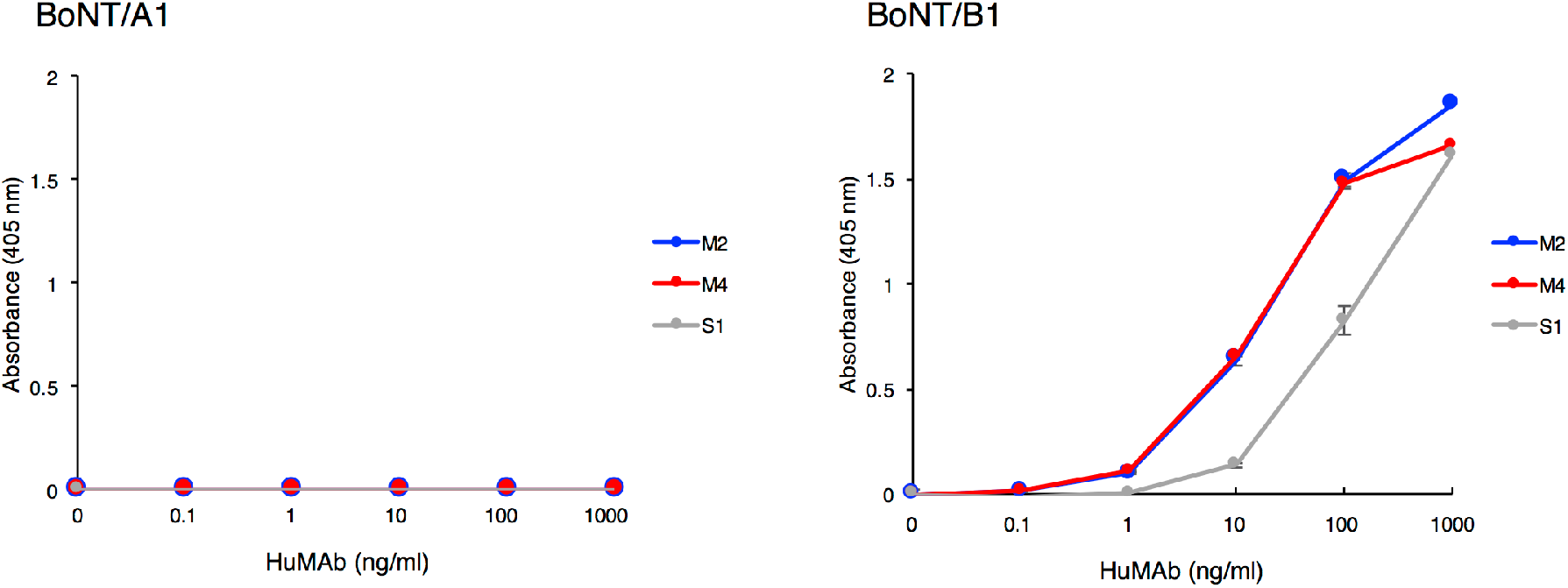
Binding of HuMAbs to BoNT/A1 and BoNT/B1. Binding of HuMAbs to BoNT was analyzed by ELISA. HuMAbs (1.0–1000 ng/ml) were added to plates coated with BoNT/A1 or BoNT/B1. After washing, bound HuMAbs were detected by anti–human IgG antibody conjugated with HRP.

**Figure 2.**
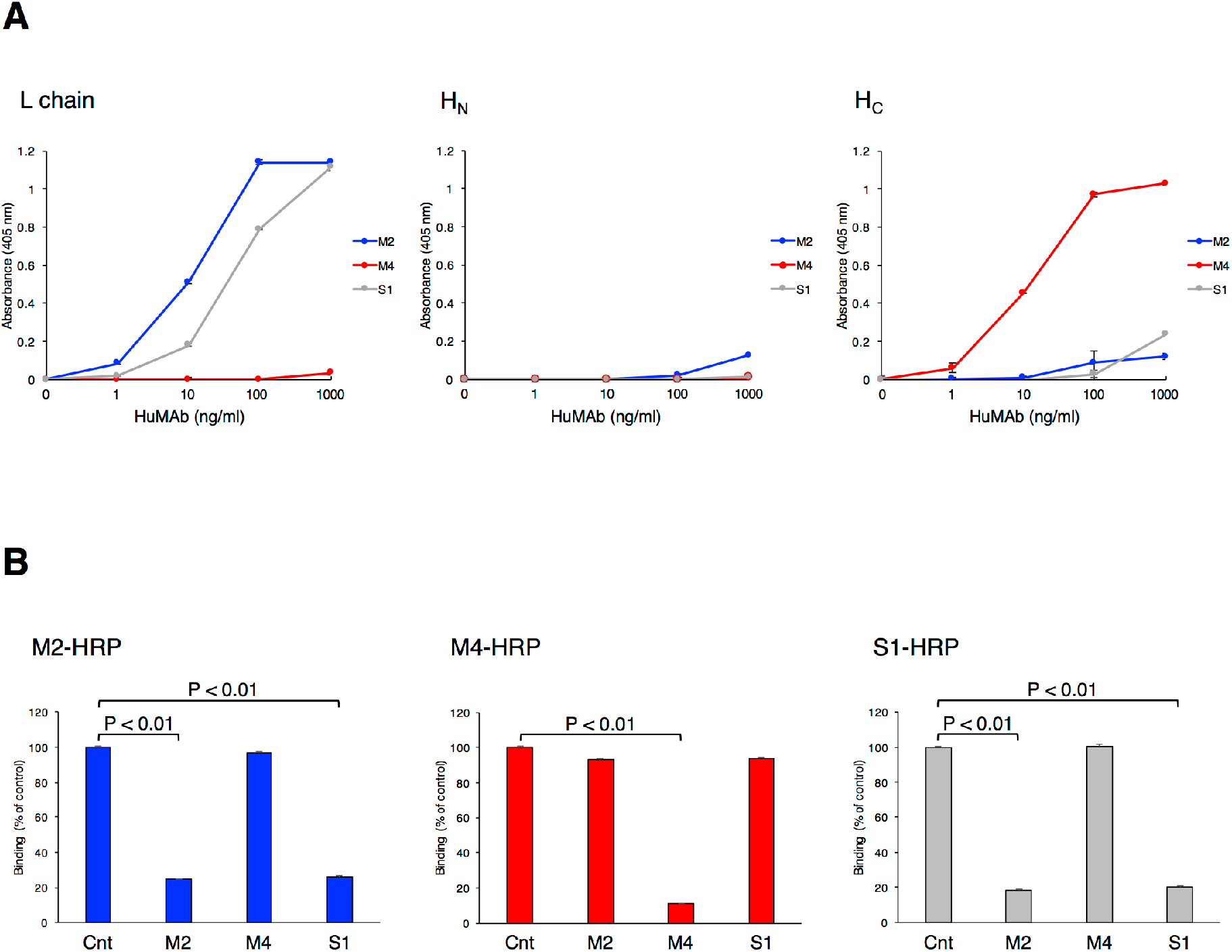
Binding domains of BoNT/B1 recognized by HuMAbs. (A) Binding of HuMAbs to recombinant L chain, H_N_, and H_C_ were analyzed by ELISA. HuMAbs (1.0–1000 ng/ml) were added to plates coated with recombinant L chain, H_N_, or H_C_. After washing, bound HuMAbs were detected by anti-human IgG antibody conjugated with HRP. (B) Binding epitopes of HuMAbs were confirmed by competitive binding ELISA. Non-labeled HuMAbs (2.5 μg/ml) were added to plates coated with BoNT/B1, and then HRP-labeled M2 (1:5000), M4 (1:5000), or S1 (1:1000) was added. Bound HRP–labeled HuMAbs were detected. Results are expressed as percent binding of each HRP-labeled HuMAb, with the 100% value defined by binding in the absence of non-labeled HuMAbs. Statistical analyses were performed with the Student’s t-test.

### Neutralization activity of HuMAbs

The neutralization activity of HuMAbs against BoNT/B1 was determined by mouse bioassay. A dose of 10 mouse intraperitoneal (i.p.) lethal dose 50% (LD_50_) of BoNT/B1 was incubated with 100 μg of HuMAb for 1 h prior to i.p. injection into mice. Control mice treated with PBS died within 12 h. By contrast, mice treated with M4 were partially protected, as evidenced by an increased time-to-death compared with control mice. Furthermore, complete protection was observed in mice treated with M2 (Fig. 3A). Because the S1-producing hybridoma had low productivity, we could not obtain a sufficient quantity of S1 to test its neutralization activity in the mouse bioassay.

**Figure 3.**
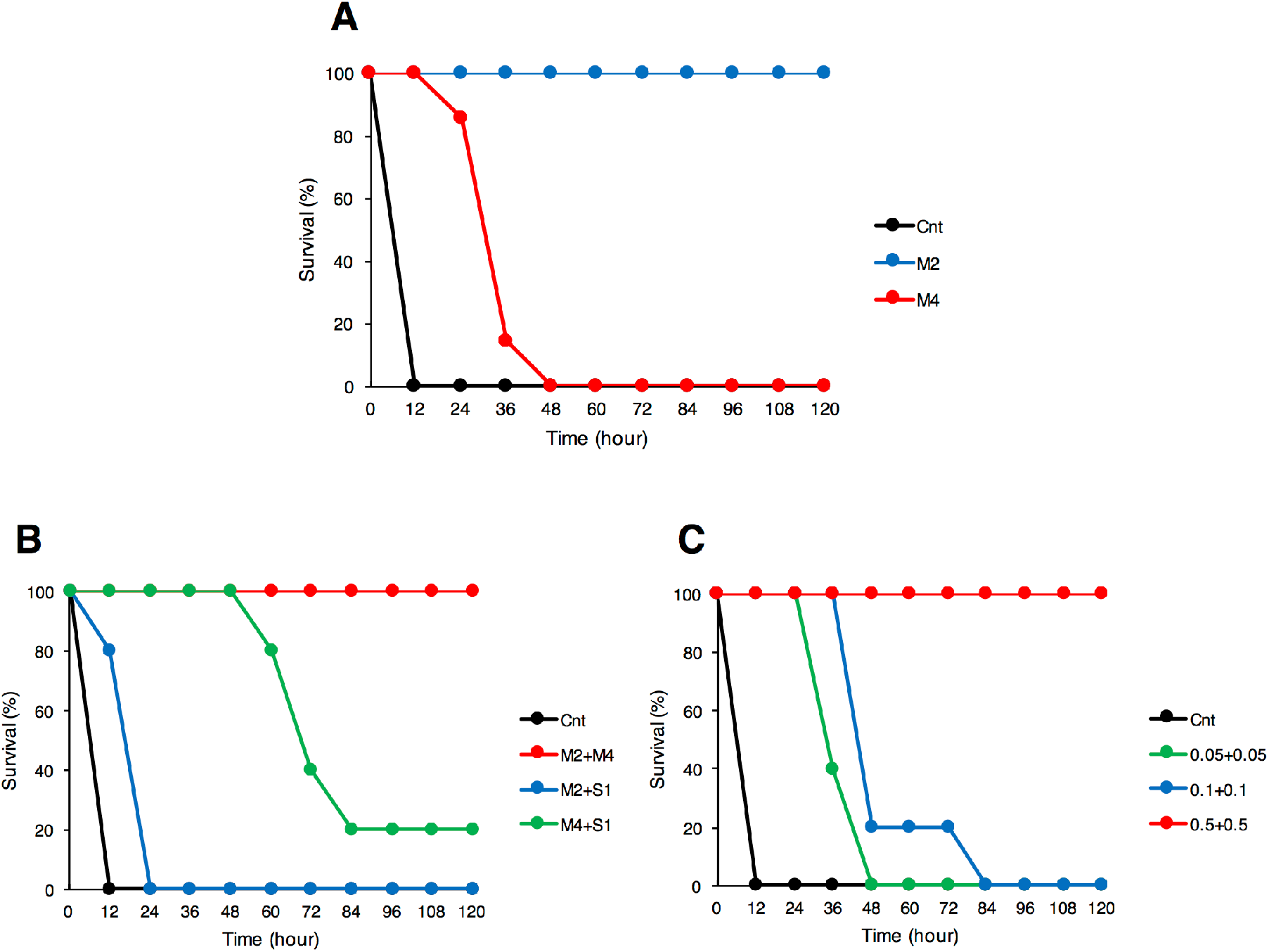
Neutralization activity of HuMAbs against BoNT/B1. The neutralization activity of HuMAbs against BoNT/B1 was determined by mouse bioassay. (A) A dose of 10 i.p. LD_50_ of BoNT/B1 was incubated with 100 μg of M2 or M4 for 1 h prior to i.p. injection into mice. Control mice (Cnt) were treated with PBS instead of HuMAbs. Mice were observed for morbidity and mortality for 2 weeks. *n* = 5 per group. (B) Neutralization of BoNT/B1 with two combinations of HuMAbs. A dose of 10 i.p. LD_50_ of BoNT/B1 was incubated with a mixture of HuMAbs (5.0 μg each) and administered by i.p. injection into mice. Control mice (Cnt) were treated with PBS instead of HuMAbs. Mice were observed for morbidity and mortality for 2 weeks. Cnt, *n* = 10 per group, M2+M4, *n* = 5 per group, M2+S1, *n* = 5 per group, M4+S1, *n* = 5 per group. (C) Neutralization of BoNT/B1 with the combination of M2 and M4. A dose of 10 i.p. LD_50_ of BoNT/B1 was incubated with a mixture of M2 and M4 (0.05, 0.1, or 0.5 μg each) and then administered by i.p. injection into mice. Control mice (Cnt) were treated with PBS instead of HuMAbs. Mice were observed for morbidity and mortality for 2 weeks. Cnt, *n* = 10 per group, 0.05 μg each, *n* = 5 per group, 0.1 μg each, *n* = 5 per group, 0.5 μg each, *n* = 5 per group.

It is known that a combination of monoclonal antibodies shows a synergistic effect neutralizing BoNT (23–26). Thus, we next tested for potential synergy of a combination of HuMAbs. A dose of 10 i.p. LD_50_ of BoNT/B1 was incubated with two combinations of HuMAbs (5.0 μg each) and injected into mice. The combination of M2 and S1 showed a very weak synergistic effect, and the combination of M4 and S1 showed partial protection. By contrast, all of the mice that received the combination of M2 and M4 (M2+M4, 5.0 μg or 0.5 μg each) survived and had no symptoms (Fig. 3B, 3C), indicating that this combination acted highly synergistically in neutralizing BoNT/B1. Based on these results, we used M2 and M4 in subsequent experiments. We obtained the sequences of the variable domains of the H and L chains of these antibodies using reverse transcription polymerase chain reaction (RT-PCR) with consensus primers (Fig. 4).

**Figure 4.**
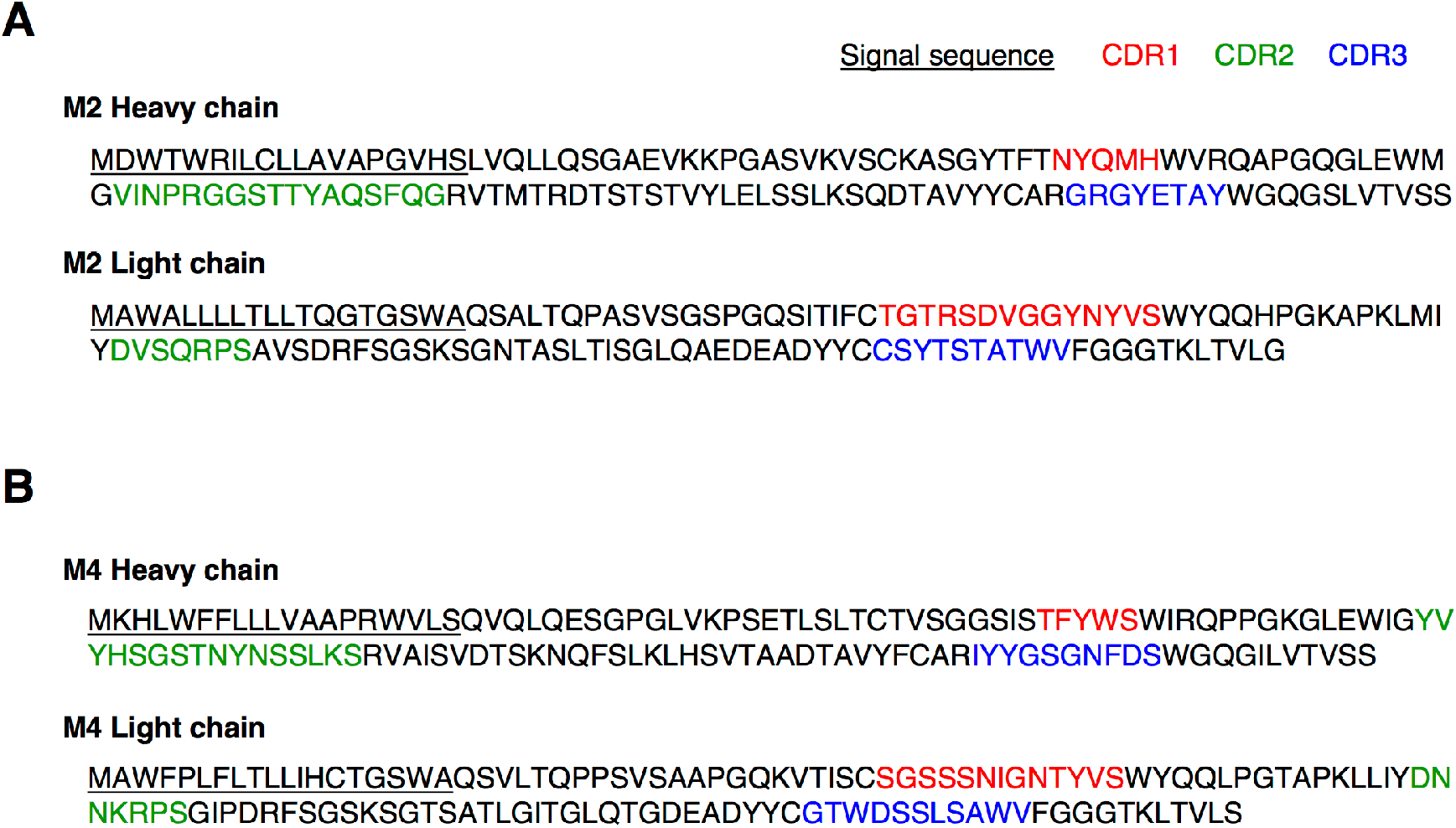
Amino acid sequences of the variable regions of the heavy and light (lambda) chains of (A) M2 and (B) M4. Complementarity-determining regions (CDRs) are indicated in red (CDR1), green (CDR2), and blue (CDR3). Signal sequences are underlined.

BoNTs are always produced along with one or more neurotoxin-associated proteins that non-covalently associate with the BoNT to form progenitor toxin complexes (PTCs). *C*. *botulinum* type B strains produce the progenitor toxins M-PTC and L-PTC (27). M-PTC is composed of BoNT and non-toxic non-HA (NTNHA), whereas L-PTC consists of BoNT, NTNHA, and haemagglutinin (HA). Food-borne botulism and infant botulism are caused by intestinal absorption of these toxin complexes. Therefore, analysis of the neutralization effect of HuMAbs using progenitor toxins is important for development of therapeutic agents for botulism. We tested the therapeutic effect of HuMAbs against orally administrated L-PTC in mice (a post-exposure treatment mouse model). The symptoms of botulism in mice, including fuzzy hair, muscle weakness, and respiratory failure, were observed at 12–24 h after oral ingestion of L-PTC, and untreated mice died within 72 h. By contrast, complete survival was observed following M2+M4 (0.5 μg each) treatment at 12 h after oral administration of L-PTC, and partial survival was observed when treatment was given at 24 and 36 h (Fig. 5A). Furthermore, we tested the preventive effect of passive immunization with HuMAbs (a pre-exposure prevention mouse model). In this model, mice were administered i.p. BoNT/B1 (10 i.p. LD_50_). Non-treated mice died within 24 h. By contrast, pre-administration with M2+M4 (0.5 μg each) up to 3 days before injection of BoNT/B1 resulted in complete protection against the lethal dose of BoNT/B1. Partial protection was observed in mice administered M2+M4 at 5 or 7 days before BoNT injection (Fig. 5B).

**Figure 5.**
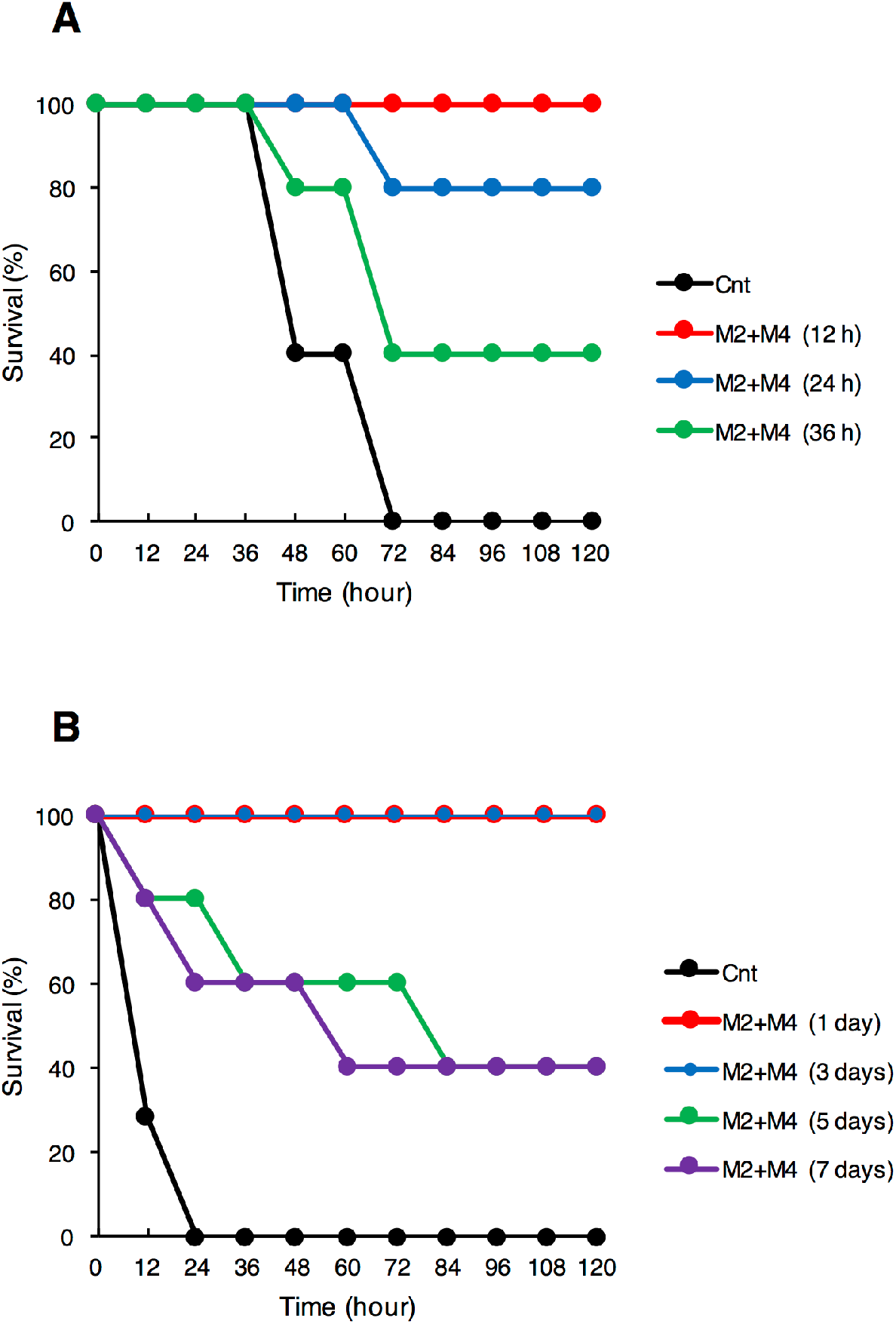
Therapeutic and preventive effects of HuMAbs (M2+M4) against BoNT/B1 intoxication. (A) Mice were orally administered progenitor toxin (L-PTC, 10 ng), and subsequently administered M2+M4 (0.5 μg each) by i.p. injection at 12, 24, or 36 h after the oral administration of L-PTC. Control mice (Cnt) were not treated with HuMAbs. Mice were observed for morbidity and mortality for 2 weeks. *n* = 5 per group. (B) Mice received i.p. injection of M2+M4 (0.5 μg each) and were then challenged at 1, 3, 5, or 7 days later with i.p. administration of 10 i.p. LD_50_ BoNT/B1. Control mice (Cnt) were not injected with HuMAbs. Mice were observed for morbidity and mortality for 2 weeks. Cnt, *n* = 7 per group, M2+M4 (1 day), *n* = 5 per group, M2+M4 (3 days), *n* = 5 per group, M2+M4 (5 days), *n* = 5 per group. M2+M4 (7 days), *n* = 5 per group.

### Binding and neutralization activity of HuMAbs against subtypes BoNT/B2 and BoNT/B6

We next tested the breadth of the neutralization activity of M2+M4 by testing their efficacy against subtypes BoNT/B2 and BoNT/B6, which caused infant botulism in Japan (28,29). ELISA showed that M2 and M4 bound strongly to BoNT/B2 and BoNT/B6 as well as BoNT/B1 (Fig. 6A). In the neutralization test, mice injected with 10 ng of BoNT/B2 or 2.5 ng of BoNT/B6 died within 12 h, demonstrating a degree of lethality that was similar to that of 10 i.p. LD_50_ of BoNT/B1. M2+M4 (0.5 μg each) completely neutralized BoNT/B2 and BoNT/B6 (Fig. 6B).

**Figure 6.**
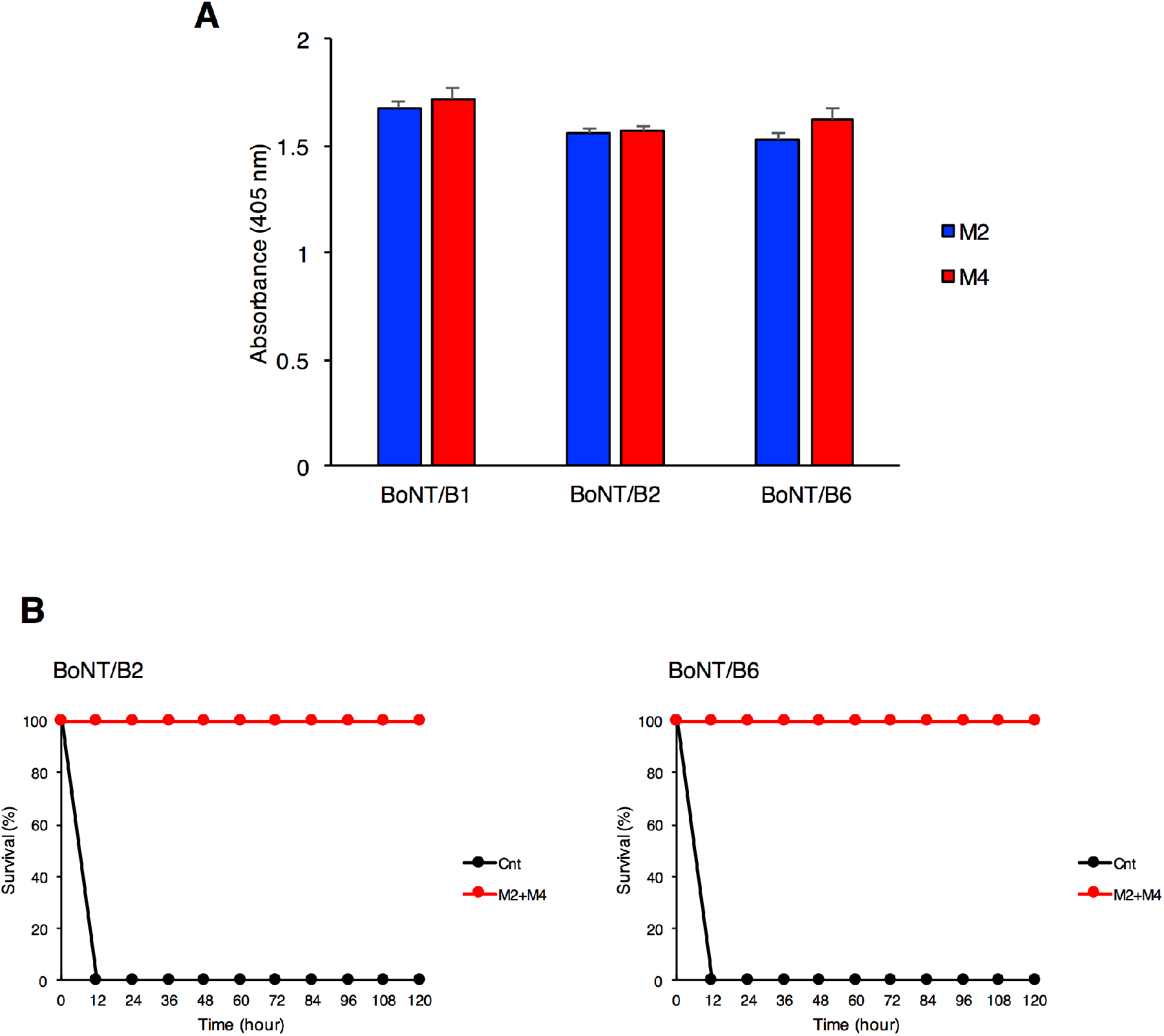
Binding and neutralization activity of HuMAbs against BoNT/B2 and BoNT/B6. (A) Binding of HuMAbs to BoNT/B2 and BoNT/B6 were analyzed by ELISA. HuMAbs (0.1 μg/ml) were added to plates coated with BoNT/B2 or BoNT/B6. After washing, bound HuMAbs were detected by anti–human IgG antibody conjugated with HRP. (B) Doses of 10 ng of BoNT/B2 or 2.5 ng of BoNT/B6 were incubated with M2+M4 (0.5 μg each) and administered by i.p. injection into mice. Control mice (Cnt) were treated with PBS instead of HuMAbs. Mice were observed for morbidity and mortality for 2 weeks. *n* = 5 per group.

## DISCUSSION

Mouse or humanized monoclonal antibodies against BoNT/B have been reported in many studies (25,30–34), but there is very little information on fully human monoclonal antibodies. HuMAbs have been shown to effectively neutralize BoNT/B (35), and when used therapeutically in humans, are considered to carry a lower risk of side effects such as serum sickness and anaphylaxis than mouse, humanized, or equine antibodies. Thus, HuMAb therapy may be an effective treatment for human botulism. In this study, we developed fully human monoclonal antibodies specific for BoNT/B1 using the SPYMEG cell line. Among these HuMAbs, M2 specifically bound to the L chain of BoNT/B1 and showed potent neutralization activity in a mouse bioassay (approximately ≥ 100 i.p. LD_50_/mg of antibody) (Fig. 3A). Mechanistically, M2 may directly inhibit the proteolytic activity of the L chain or prevent the translocation of the L chain into the cytosol. In fact, both of these mechanisms were reported in a HuMAb that recognized the L chain of BoNT/A (24). On the other hand, M4 bound to the H_C_ (receptor binding domain) and showed partial neutralization activity. M4 may prevent the binding of BoNT/B1 to neuronal cells (25). Meanwhile, the reactivity of M4 with BoNT/B1 was very weak in western blot analysis (denaturing condition) (data not shown). Hence, M4 may recognize the conformational epitope of BoNT/B1. The epitope binding details and neutralization mechanisms of M2 and M4 are currently being analyzed.

Several studies have confirmed a synergistic effect when two or more antibodies are combined (23–26). The combination of the HuMAbs M2 and M4, which recognize non-overlapping epitopes, was able to completely neutralize BoNT/B1 with a potency greater than 10,000 i.p. LD_50_/mg of antibodies (Fig. 3B, C). This potency is over 12.5 times greater than that of BabyBIG (18) (800 LD_50_/mg of antibodies), an orphan drug that consists of human-derived anti-botulinum toxin antibodies. By contrast, the combination of M2 and S1, which recognize overlapping epitopes, showed no synergistic effect. This result suggests that the recognition of non-overlapping epitopes is critical for a synergistic effect in BoNT neutralization. Meanwhile, the combination of M4 and S1 showed only partial protection even though M4 and S1 recognize non-overlapping epitopes. The antigen binding affinity of S1 might be lower than that of M2, or the amino acid sequence of BoNT/B1 recognized by S1 might be slightly different from that of M2, both of which are factors that may have influenced the synergistic effect of M4+S1.

Food-borne botulism and infant botulism, which account for the majority of cases of human botulism, are caused by the intestinal absorption of the progenitor toxins M-PTC and L-PTC. The oral toxicity of L-PTC is much higher than the toxicity of M-PTC and BoNT individually (10,36). Thus, L-PTC is considered to be the predominant contributor to the onset of food-borne botulism and presumably infant botulism. In human food-borne botulism, the signs and symptoms of botulism typically begin between 12 and 36 h after ingestion of the toxin (9). Therefore, we used a dose of L-PTC at which almost all control mice presented with botulism symptoms approximately 12–24 h after oral administration of L-PTC. In this mouse model, M2+M4 provided complete protection when administered within 12 h of oral ingestion of L-PTC, and partial survival was observed at 24 and 36 h after L-PTC ingestion. Our results show that M2+M4 exerts potent protective activity even after botulism symptoms have developed, but early administration is important. In fact, in the case of food-borne botulism, it is recommended to administer antitoxin as early as possible after toxin exposure. Good treatment outcomes are directly correlated with early administration of antitoxins (9). Preventive countermeasures against BoNT are also needed for individuals at risk of BoNT exposure, such as first responders in contaminated areas (for example, in instances of bioterrorism or outbreak). Previous studies have reported that passive immunization with HuMAbs protects against the onset of type A botulism (37,38). In this study, M2+M4 showed a long-term preventive effect against a lethal dose of BoNT/B1 (Fig. 5B). It is presumed that because of their low immunogenicity and high affinity to human neonatal Fc receptor compared with mouse antibodies, the HuMAbs M2 and M4 may exist longer in the human bloodstream and therefore neutralize BoNT/B1 more effectively and for a longer period of time (39). Taken together, these data indicate that M2+M4 has sufficient therapeutic and preventive effects against botulism to permit practical use. Fully human monoclonal antibodies showing therapeutic and preventive effects against type B botulism in mouse models have never been reported before.

We also examined the neutralization effect of M2+M4 against two other BoNT/B subtypes, namely BoNT/B2 and BoNT/B6. For BoNT/B, currently eight subtypes, B1 to B8 are known, with most BoNT/B strain producing B1 and B2 (40–42). The BoNT/B1-producing strain Okra was isolated from a case of food-borne botulism (43). The BoNT/B2-producing strain 111 and the BoNT/B6-producing strain Osaka05 were isolated from infant botulism cases in Japan (28,29). BoNT/B2 and BoNT/B6 show different antigenic and biological properties than BoNT/B1 (28,29,44,45). However, M2 and M4 exhibited strong binding and high neutralization activity against BoNT/B2 and BoNT/B6 as well as against BoNT/B1 (Fig. 6). These data indicate that M2 and M4 may be effective for the treatment and prevention of botulism caused by several subtypes of BoNT/B.

In conclusion, we developed anti-BoNT/B antibody–producing hybridomas using the SPYMEG cell line, and obtained two neutralizing antibodies, M2 and M4. This method can be applied to other botulinum serotypes, and it will be possible to maintain a stable supply of HuMAbs using these hybridomas. Because of their human origin, M2 and M4 are safe for the treatment of human botulism, including infant botulism. Furthermore, we found that the combination of M2 and M4 had potent neutralization activity, greater than that of BabyBIG, against BoNT/B1, and neutralized two other subtypes. Additionally, M2+M4 showed therapeutic and preventive effects against botulism in mouse models. These data indicate that the fully human monoclonal antibodies, M2 and M4, are promising candidates for the development of human therapeutics and prophylactics for BoNT/B intoxication.

## MATERIALS AND METHODS

### Ethics statement

Human materials were collected using protocols approved by the Institutional Review Board of the Osaka University Research Institute for Microbial Diseases (#20-8). Written informed consent was obtained from the participants. Animal studies were conducted under the applicable laws and guidelines for the care and use of laboratory animals at the Osaka University Research Institute for Microbial Diseases and Kanazawa University. They were approved by the Animal Experiment Committee of the Osaka University Research Institute for Microbial Diseases (#H21-27-0, #H27-02-0) and Kanazawa University (AP-163710).

### Preparation of BoNT/A and BoNT/B

*C. botulinum* serotype B1 strain Okra was cultured using a cellophane tube procedure, and the culture supernatant was obtained. Progenitor toxins (M-PTC and L-PTC) were purified from culture supernatant, and BoNT/B1 and a non-toxic component were prepared from L-PTC as described previously (46). The lethal toxicity of BoNT/B1 was determined using mouse bioassay (47,48). BoNT/A1 (strain 62A), BoNT/B2 (strain 111), and BoNT/B6 (strain Osaka05) were provided by Dr. T. Kohda (Osaka Prefecture University).

### Inoculation of botulinum toxoid vaccine

Two healthy adult volunteers were inoculated four or five times with a tetravalent botulinum toxoid vaccine (types A, B, E, and F) (22) provided by Dr. M. Takahashi (National Institute of Infectious Diseases). The toxoid was injected intramuscularly in 0.5-ml doses. Blood samples were collected from each volunteer and the plasma antibody titers against BoNT/A1 and BoNT/B1 were tested by ELISA.

### Preparation of HuMAbs

Peripheral blood samples (10 ml) were collected from volunteers 9 or 18 days after the last immunization, and peripheral blood mononuclear cells (PBMCs) were purified by Ficoll (GE Healthcare, Buckinghamshire, UK) gradient centrifugation. The PBMCs were fused with cells from the mouse–human fusion partner cell line SPYMEG (Medical & Biological Laboratories, Nagoya, Japan) at a ratio of 10:1 with polyethylene glycol (Roche, Basel, Switzerland). Fused cells were cultured in Dulbecco’s modified Eagle medium (DMEM, Thermo Fisher Scientific, Waltham, MA, USA) supplemented with 15% heat-inactivated fetal bovine serum in 96-well plates for 10–14 days in the presence of hypoxanthine–aminopterin–thymidine (HAT, Thermo Fisher Scientific). The first screening of the culture medium for antibodies specific to BoNT/A1 or BoNT/B1 was performed by ELISA. BoNT/A1 or BoNT/B1-specific antibody-positive wells were next subjected to cell cloning by limiting dilution. The second screening was also performed by ELISA. Each stable hybridoma was cultured in serum-free medium (Thermo Fisher Scientific), and then the supernatant was collected. Isotypes of HuMAbs in culture supernatant were determined by western blot using anti–human IgG conjugated with horseradish peroxidase (HRP) (Bio-Rad Laboratories, Berkeley, CA, USA, cat. 172–1050), anti–human IgM conjugated with HRP (Invitrogen, Carlsbad, CA, USA, cat. 627520), or anti-human IgA conjugated with HRP (Invitrogen, cat. AHI0104). IgG antibodies were purified from the supernatant using a Protein G column (GE Healthcare). The subclass of each IgG antibody was determined using a human IgG subclass ELISA kit (Invitrogen).

### Preparation of recombinant proteins

Purified DNA from *C. botulinum* type B strain Okra was used as a template for the amplification of DNA encoding L chain (aa 1-430), H_N_ (aa 449-858), and H_C_ (aa 858-1291) by PCR. The amplified DNAs were inserted into *Nco*I-*Xho*I site of the pET28b vector (Novagen, Merck Millipore, Madison, WI, USA). Recombinant proteins were expressed as C-terminally His-tagged proteins in *E. coli* strain BL21-CodonPlus (DE3)-RIL (Agilent Technologies, Santa Clara, CA, USA), and purified using HisTrap HP (GE Healthcare).

### Binding assay using ELISA

The 96-well plates (Corning, Corning, NY, USA) were coated with BoNT, recombinant L chain, recombinant H_N_, or recombinant H_C_ (300 ng/well) for 2 h at 37 °C. The wells were washed with phosphate-buffered saline (PBS) containing 0.05% Tween20 (Sigma-Aldrich, St. Louis, MO, USA) (PBS-T) and blocked with 0.2% bovine serum albumin (BSA, Sigma-Aldrich)/PBS-T overnight at 4 °C. Human plasma samples or HuMAbs were added to a well and incubated for 2 h at 37 °C. After washing, anti–human IgG conjugated with HRP (Bio-Rad Laboratories, cat. 172-1050, or Jackson ImmunoResearch, West Grove, PA, USA, cat. 309-035-003) was added and incubated for 2 h at 37 °C. Plates were washed again and then incubated with substrate solution (*o*-phenylendiamine, Nacalai Tesque, Kyoto, Japan or ABTS, Roche) for 20 min at 37 °C, and absorbance values at Abs_492_ or Abs_405_ were measured, respectively.

### Competition ELISA

M2, M4, and S1 were labeled with HRP using Perioxidase Labeling Kits (Dojindo Molecular Technologies, Kumamoto, Japan). The optimal dilutions of HRP-labeled HuMAbs were determined by ELISA based on an Abs_405_ value around 2.0. Non-labeled HuMAb (2.5 μg/ml) was added to plates coated with BoNT/B1 and incubated for 2 h at 37 °C. After washing, HRP-labeled M2 (1:5000), M4 (1:5000), or S1 (1:1000) was added and incubated for 2 h at 37 °C. Plates were washed again and incubated with substrate solution (ABTS) for 20 min at 37 °C, and absorbance values at Abs405 were measured.

### Sequencing of HuMAb variable region gene segments

Total RNA was extracted from the hybridomas using an RNeasy Mini Kit (Qiagen, Hilden, Germany), and cDNA was synthesized by RT-PCR using a SuperScript^®^ VILO^TM^ cDNA Synthesis Kit (Invitrogen). The coding regions of the H and L chains of M2 and M4 were amplified by PCR using KOD-Plus-Neo (TOYOBO, Osaka, Japan), with the following primers: 5’-ATGGACTGGACCTGGAGGATCCTC-3’ (M2 H chain sense primer), 5’-ATGAAACACCTGTGGTTCTTCCTCCT-3’ (M4 H chain sense primer), and 5’-CTCCCGCGGCTTTGTCTTGGCATTA-3’ (H chain antisense primer); and 5’-ATGSCCTGGGCTCYKCTSCTCCTS-3’ (M2 L chain sense primer), 5’-ATGGCCTGGWYYCCTCTCYTYCTS-3’ (M4 L chain sense primer), and 5’-TGGCAGCTGTAGCTTCTGTGGGACT-3’ (L chain antisense primer).

### Neutralization test (mouse bioassay)

The neutralization activity of HuMAbs was tested by mouse bioassay. Female ddY mice were purchased from SLC (Shizuoka, Japan) and were used at 4 weeks of age. One or more HuMAbs was incubated with BoNT/B1, BoNT/B2, or BoNT/B6 at room temperature for 1 h prior to i.p. injection (in a total volume of 500 μl) into mice. In control mice, BoNT was incubated with PBS instead of HuMAbs. In the botulism treatment (post-exposure) model, the B1 subtype progenitor toxin (L-PTC, 10 ng in a volume of 300 μl) was orally administered and M2+M4 was subsequently administered at 12, 24, and 36 h after oral administration of L-PTC by i.p. injection. In the botulism prevention (pre-exposure) model, M2+M4 was administered by i.p. injection and mice were then challenged at 1, 3, 5, or 7 days later with 10 i.p. LD_50_ BoNT/B1 by i.p. injection. Mice were observed for morbidity and mortality for 2 weeks.

## Abbreviations

aa: amino acid
BoNT: botulinum neurotoxin
BSA: bovine serum albumin
DMEM: Dulbecco’s modified Eagle medium
ELISA: enzyme linked immunosorbent assay
HA: haemagglutinin
HAT: hypoxanthine–aminopterin–thymidine
H_C_: C-terminal of heavy chain
H chain: heavy chain
H_N_: N-terminal of heavy chain
HRP: horseradish peroxidase
HuMAb: human monoclonal antibody
i.p.: intraperitoneal
L chain: light chain
LD_50_: lethal dose 50%
NTNHA: non-toxic non-HA
PBMC: peripheral blood mononuclear cell
PBS: phosphate buffered saline
PTC: progenitor toxin complex
RT-PCR: reverse transcription polymerase chain reaction
SNAP-25: Synaptosomal associated protein of 25 kDa
SNARE: soluble *N*-ethylmaleimide-sensitive fusion protein attachment protein receptor
SV2: synaptic vesicle protein 2
VAMP: vesicle-associated membrane protein

## ACKNOWLEDGMENTS

We would like to thank Dr. Y. Sugawara, A. Sano, K. Sasaki, and C. Aoki (Research Institute for Microbial Diseases, Osaka University, Japan); Dr. K. Hosomi (Department of Veterinary Sciences, School of Life and Environmental Sciences, Osaka Prefecture University, Japan); and S. Akagi, Y. Koino, and K. Imazaki (Department of Bacteriology, Graduate School of Medical Sciences, Kanazawa University, Japan) for their technical assistance. We also thank Dr. T. Kenri, Dr. A. Yamamoto, and Dr. M. Iwaki (Department of Bacteriology II, National Institute of Infectious Diseases, Japan) for advice on the mouse bioassay. Finally, we thank Dr. S. Kozaki (Department of Veterinary Sciences, School of Life and Environmental Sciences, Osaka Prefecture University, Japan) for advice on botulinum toxins, Dr. M. Takahashi (Department of Bacteriology II, National Institute of Infectious Diseases, Japan) for providing the toxoid vaccine, and Dr. M. Yasugi and Dr. R. Kubota-Koketsu (Research Institute for Microbial Diseases, Osaka University, Japan) for advice on the SPYMEG cell line and cell fusion.

This work was supported by the Japan Science and Technology Agency/Japan International Cooperation Agency, Science and Technology Research Partnership for Sustainable Development, 08080924 (JST/JICA, SATREPS) (http://www.jst.go.jp/global/english/kadai/h2011_thailand.html), the Japan Agency for Medical Research and Development, AMED (Grant No. 19fk0108101h0501), and the Japan Society for the Promotion of Science KAKENHI (Grant No. 16K19123, 18K07107 and No. 18H02654).

T.M. and Y.F. designed the research and analyzed the data. T.M. performed the majority of the experiments and the analysis. S.A. provided recombinant proteins. R.M. and K.F., analyzed the sequencing of HuMAbs. M.Y. analyzed the data. A.D. and K.I. provided cells from the SPYMEG cell line and performed the cell fusion experiments. T.K. provided botulinum toxins. T.M. and Y.F. co-wrote the manuscript. All authors participated in discussion of the data and in production of the final version of the manuscript.

The authors declare no competing financial interests.

## REFERENCES

1. Schiavo G, Matteoli M, Montecucco C. 2000. Neurotoxins affecting neuroexocytosis. Physiol Rev 80: 717–766; PMID:10747206; https://doi.org/10.1152/physrev.2000.80.2.717

2. Rossetto O, Pirazzini M, Montecucco C. 2014. Botulinum neurotoxins: genetic, structural and mechanistic insights. Nat Rev microbial 12:535–549; PMID:24975322; https://doi.org/10.1038/nrmicro3295

3. Froude JW, Stiles B, Pelat T, Thullier P. 2011. Antibodies for biodefense. mAbs 3: 517–527; PMID:22123065; https://doi.org/10.4161/mabs.3.6.17621

4. Barash JR, Arnon SS. 2014. A novel strain of Clostridium botulinum that produces type B and type H botulinum toxins. J Infect Dis 209:183–91; PMID:24106296; https://doi.org/10.1093/infdis/jit449

5. Dover N, Barash JR, Hill KK, Xie G, Arnon SS. 2014. Molecular characterization of a novel botulinum neurotoxin type H gene. J Infect Dis 209: 192–202; PMID:24106295; https://doi.org/10.1093/infdis/jit450

6. Maslanka SE, Lúquez C, Dykes JK, Tepp WH, Pier CL, Pellett S, Raphael BH, Kalb SR, Barr JR, Rao A, Johnson EA. 2016. A Novel Botulinum Neurotoxin, Previously Reported as Serotype H, Has a Hybrid-Like Structure With Regions of Similarity to the Structures of Serotypes A and F and Is Neutralized With Serotype A Antitoxin. J Infect Dis 213: 379–85; PMID:26068781; https://doi.org/10.1093/infdis/jiv327

7. Fan Y, Barash JR, Lou J, Conrad F, Marks JD, Arnon SS. 2016. Immunological Characterization and Neutralizing Ability of Monoclonal Antibodies Directed Against Botulinum Neurotoxin Type H. J Infect Dis 213:1606–14; PMID:26936913; https://doi.org/10.1093/infdis/jiv770

8. Yao G, Lam KH, Perry K, Weisemann J, Rummel A, Jin R. 2017. Crystal Structure of the Receptor-Binding Domain of Botulinum Neurotoxin Type HA, Also Known as Type FA or H. Toxins (Basel) 9:93; PMID:28282873; https://doi.org/10.3390/toxins9030093

9. Arnon SS, Schechter R, Inglesby TV, Henderson DA, Bartlett JG, Ascher MS, Eitzen E, Fine AD, Hauer J, Layton M, Lillibridge S, Osterholm MT, O’Toole T, Parker G, Perl TM, Russell PK, Swerdlow DL, Tonat K; Working Group on Civilian Biodefense. 2001. Botulinum toxin as a biological weapon: medical and public health management. JAMA 285: 1059–1070; PMID:11209178; http://dx.doi.org/10.1001/jama.285.8.1059

10. Matsumura T, Sugawara Y, Yutani M, Amatsu S, Yagita H, Kohda T, Fukuoka S, Nakamura Y, Fukuda S, Hase K, Ohno H, Fujinaga Y. 2015. Botulinum toxin A complex exploits intestinal M cells to enter the host and exert neurotoxicity. Nat Commun 6: 6255; PMID:25687350; https://doi.org/10.1038/ncomms7255

11. Dong M, Yeh F, Tepp WH, Dean C, Johnson EA, Janz R, Chapman ER. 2006. SV2 is the protein receptor for botulinum neurotoxin A. Science 312: 592–596; PMID:16543415; https://doi.org/10.1126/science.1123654

12. Dong M, Liu H, Tepp WH, Johnson EA, Janz R, Chapman ER. 2008. Glycosylated SV2A and SV2B mediate the entry of botulinum neurotoxin E into neurons. Mol Biol Cell 19: 5226–5237; PMID:18815274; https://doi.org/10.1091/mbc.e08-07-0765

13. Fu Z, Chen C, Barbieri JT, Kim JJ, Baldwin MR. 2009. Glycosylated SV2 and gangliosides as dual receptors for botulinum neurotoxin serotype F. Biochemistry 48: 5631–5641; PMID:19476346; https://doi.org/10.1021/bi9002138

14. Nishiki T, Kamata Y, Nemoto Y, Omori A, Ito T, Takahashi M, Kozaki S. 1994. Identification of protein receptor for Clostridium botulinum type B neurotoxin in rat brain synaptosomes. J Biol Chem 269: 10498–10503; PMID:8144634;

15. Nishiki T, Tokuyama Y, Kamata Y, Nemoto Y, Yoshida A, Sato K, Sekiguchi M, Takahashi M, Kozaki S. 1996. The high-affinity binding of Clostridium botulinum type B neurotoxin to synaptotagmin II associated with gangliosides GT1b/GD1a. FEBS Lett 378: 253–257; PMID:8557112; https://doi.org/10.1016/0014-5793(95)01471-3

16. Hoch DH, Romero-Mira M, Ehrlich BE, Finkelstein A, DasGupta BR, Simpson LL. 1985. Channels formed by botulinum, tetanus, and diphtheria toxins in planar lipid bilayers: relevance to translocation of proteins across membranes. Proc Natl Acad Sci U S A 82: 1692–1696; PMID:3856850; https://doi.org/10.1073/pnas.82.6.1692

17. Koriazova LK, Montal M. 2003. Translocation of botulinum neurotoxin light chain protease through the heavy chain channel. Nat Struct Biol 10:13–8; PMID:12459720; https://doi.org/10.1038/nsb879

18. Arnon SS, Schechter R, Maslanka SE, Jewell NP, Hatheway CL. 2006. Human botulism immune globulin for the treatment of infant botulism. N Engl J Med 354: 462–471; PMID:16452558; https://doi.org/10.1056/NEJMoa051926

19. Kubota-Koketsu R, Mizuta H, Oshita M, Ideno S, Yunoki M, Kuhara M, Yamamoto N, Okuno Y, Ikuta K. 2009. Broad neutralizing human monoclonal antibodies against influenza virus from vaccinated healthy donors. Biochem Biophys Res Commun 387:180–185; PMID:19580789; https://doi.org/10.1016/j.bbrc.2009.06.151

20. Pan Y, Sasaki T, Kubota-Koketsu R, Inoue Y, Yasugi M, Yamashita A, Ramadhany R, Arai Y, Du A, Boonsathorn N, Ibrahim MS, Daidoji T, Nakaya T, Ono K, Okuno Y, Ikuta K, Watanabe Y. 2014. Human monoclonal antibodies derived from a patient infected with 2009 pandemic influenza A virus broadly cross-neutralize group 1 influenza viruses. Biochem Biophys Res Commun 450: 42–48; PMID:24858683; https://doi.org/10.1016/j.bbrc.2014.05.060

21. Misaki R, Fukura N, Kajiura H, Yasugi M, Kubota-Koketsu R, Sasaki T, Momota M, Ono K, Ohashi T, Ikuta K, Fujiyama K. 2016. Recombinant production and characterization of human anti-influenza virus monoclonal antibodies identified from hybridomas fused with human lymphocytes. Biologicals 44:394–402; PMID:27464991; https://doi.org/10.1016/j.biologicals.2016.05.006

22. Torii Y, Tokumaru Y, Kawaguchi S, Izumi N, Maruyama S, Mukamoto M, Kozaki S, Takahashi M. 2002. Production and immunogenic efficacy of botulinum tetravalent (A, B, E, F) toxoid. Vaccine 20: 2556–61; PMID:12057613; https://doi.org/10.1016/S0264-410X(02)00157-3

23. Nowakowski A, Wang C, Powers DB, Amersdorfer P, Smith TJ, Montgomery VA, Sheridan R, Blake R, Smith LA, Marks JD. 2002. Potent neutralization of botulinum neurotoxin by recombinant oligoclonal antibody. Proc Natl Acad Sci U S A 99: 11346–11350; PMID:12177434; https://doi.org/10.1073/pnas.172229899

24. Adekar SP, Takahashi T, Jones RM, Al-Saleem FH, Ancharski DM, Root MJ, Kapadnis BP, Simpson LL, Dessain SK. 2008. Neutralization of botulinum neurotoxin by a human monoclonal antibody specific for the catalytic light chain. PLoS One 3: e3023; PMID:18714390; https://doi.org/10.1371/journal.pone.0003023

25. Chen C, Wang S, Wang H, Mao X, Zhang T, Ji G, Shi X, Xia T, Lu W, Zhang D, Dai J, Guo Y. 2012. Potent neutralization of botulinum neurotoxin/B by synergistic action of antibodies recognizing protein and ganglioside receptor binding domain. PLoS One 7:e43845; PMID:22952786; https://doi.org/10.1371/journal.pone.0043845

26. Diamant E, Lachmi BE, Keren A, Barnea A, Marcus H, Cohen S, David AB, Zichel R. 2014. Evaluating the synergistic neutralizing effect of anti-botulinum oligoclonal antibody preparations. PLoSOne 9:e87089;PMID:24475231; https://doi.org/10.1371/journal.pone.0087089

27. Fujinaga Y, Sugawara Y, Matsumura T. 2013. Uptake of botulinum neurotoxin in the intestine. Curr Top Microbiol Immunol 364: 45–59; PMID:23239348; https://doi.org/10.1007/978-3-642-33570-9_3.

28. Kozaki S, Kamata Y, Nishiki T, Kakinuma H, Maruyama H, Takahashi H, Karasawa T, Yamakawa K, Nakamura S. 1998. Characterization of Clostridium botulinum type B neurotoxin associated with infant botulism in japan. Infect Immun 66: 4811–6; PMID:9746583;

29. Umeda K, Seto Y, Kohda T, Mukamoto M, Kozaki S. 2009. Genetic characterization of Clostridium botulinum associated with type B infant botulism in Japan. J Clin Microbiol 47: 2720–8; PMID:19571018; https://doi.org/10.1128/JCM.00077-09

30. Yang GH, Kim KS, Kim HW, Jeong ST, Huh GH, Kim JC, Jung HH. 2004. Isolation and characterization of a neutralizing antibody specific to internalization domain of Clostridium botulinum neurotoxin type B. Toxicon 44:19–25; PMID:15225558; https://doi.org/10.1016/j.toxicon.2004.03.016

31. Cheng LW, Henderson TD 2nd, Lam TI, Stanker LH. 2015. Use of Monoclonal Antibodies in the Sensitive Detection and Neutralization of Botulinum Neurotoxin Serotype B. Toxins (Basel) 7: 5068–5078; PMID:26633496; https://doi.org/10.3390/toxins7124863

32. Rasetti-Escargueil C, Avril A, Chahboun S, Tierney R, Bak N, Miethe S, Mazuet C, Popoff MR, Thullier P, Hust M, Pelat T, Sesardic D. 2015. Development of human-like scFv-Fc antibodies neutralizing Botulinum toxin serotype B. MAbs 37: 1161–1177; PMID:26381852; https://doi.org/10.1080/19420862.2015.1082016

33. Fan Y, Dong J, Lou J, Wen W, Conrad F, Geren IN, Garcia-Rodriguez C, Smith TJ, Smith LA, Ho M, Pires-Alves M, Wilson BA. 2015. Monoclonal Antibodies that Inhibit the Proteolytic Activityof Botulinum NeurotoxinSerotype/B. Toxins (Basel) 7:3405–3423; PMID:26343720; https://doi.org/10.3390/toxins7093405

34. Miethe S, Mazuet C, Liu Y, Tierney R, Rasetti-Escargueil C, Avril A, Frenzel A, Thullier P, Pelat T, Urbain R, Fontayne A, Sesardic D, Hust M, Popoff MR. 2016. Development of Germline-Humanized Antibodies Neutralizing Botulinum Neurotoxin A and B. PLoS One 11:e0161446; PMID:27560688; https://doi.org/10.1371/journal.pone.0161446

35. Garcia-Rodriguez C, Geren IN, Lou J, Conrad F, Forsyth C, Wen W, Chakraborti S, Zao H, Manzanarez G, Smith TJ, Brown JL, Skerry JC, Smith LA, Marks JD. 2011. Neutralizing human monoclonal antibodies binding multiple serotypes of botulinum neurotoxin. Protein Eng Des Sel 24:321–331; PMID:21149386; https://doi.org/10.1093/protein/gzq111

36. Sakaguchi G. 1982. Clostridium botulinum toxins. Pharmacol Ther 19: 165–194; PMID:6763707; https://doi.org/10.1016/0163-7258(82)90061-4

37. Adekar SP, Segan AT, Chen C, Bermudez R, Elias MD, Selling BH, Kapadnis BP, Simpson LL, Simon PM, Dessain SK. 2011. Enhanced neutralization potency of botulinum neurotoxin antibodies using a red blood cell-targeting fusion protein. PLoS One 6:e17491; https://doi.org/10.1371/journal.pone.0017491

38. Sharma R, Zhao H, Al-Saleem FH, Ubaid AS, Puligedda RD, Segan AT, Lindorfer MA, Bermudez R, Elias M, Adekar SP, Simpson LL, Taylor RP, Dessain SK. 2014. Mechanisms of enhanced neutralization of botulinum neurotoxin by monoclonal antibodies conjugated to antibodies specific for the erythrocyte complement receptor. Mol Immunol 57: 247–254; PMID:24184879; https://doi.org/10.1016/j.molimm.2013.09.005

39. Suzuki T, Ishii-Watabe A, Tada M, Kobayashi T, Kanayasu-Toyoda T, Kawanishi T, Yamaguchi T. 2010. Importance of neonatal FcR in regulating the serum half-life of therapeutic proteins containing the Fc domain of human IgG1: a comparative study of the affinity of monoclonal antibodies and Fc-fusion proteins to human neonatal FcR. J Immunol 184: 1968–1976; PMID:20083659; https://doi.org/10.4049/jimmunol.0903296

40. Hill KK, Smith TJ, Helma CH, Ticknor LO, Foley BT, Smith TJ. 2007. Genetic diversity among Botulinum Neurotoxin-producing clostridial strains. J Bacteriol 189: 818–832; PMID:26368006; https://doi.org/10.1016/j.toxicon.2015.09.011

41. Kalb SR, Baudys J, Rees JC, Smith TJ, Smith LA, Helma CH, Hill K, Kull S, Kirchner S, Dorner MB, Pirkle JL, Barr JR. 2012. De novo subtype and strain identification of botulinum neurotoxin type B through toxin proteomics. Anal Bioanal Chem 403: 215–226; PMID:22395449; https://doi.org/10.1007/s00216-012-5767-3

42. Peck MW, Smith TJ, Anniballi F, Austin JW, Bano L, Bradshaw M, Cuervo P, Cheng LW, Derman Y, Dorner BG, Fisher A, Hill KK, Kalb SR, Korkeala H, Lindström M, Lista F, Lúquez C, Mazuet C, Pirazzini M, Popoff MR, Rossetto O, Rummel A, Sesardic D, Singh BR, Stringer SC. 2017. Historical Perspectives and Guidelines for Botulinum Neurotoxin Subtype Nomenclature. Toxins (Basel). 18; 9 PMID:28106761; https://doi.org/10.3390/toxins9010038

43. Beers WH, Reich E. 1969. Isolation and characterization of Clostridium botulinum type B toxin. J Biol Chem 244: 4473–4479; PMID:4896752;

44. Kohda T, Ihara H, Seto Y, Tsutsuki H, Mukamoto M, Kozaki S. 2007. Differential contribution of the residues in C-terminal half of the heavy chain of botulinum neurotoxin type B to its binding to the ganglioside GT1b and the synaptotagmin 2/GT1b complex. Microb Pathog 42: 72–9; PMID:17188834; https://doi.org/10.1016/j.micpath.2006.10.006

45. Kohda T, Nakamura K, Hosomi K, Torii Y, Kozaki S, Mukamoto M. 2017. Characterization of the functional activity of botulinum neurotoxin subtype B6. Microbiol Immunol 61: 482–489; PMID:28898517; https://doi.org/10.1111/1348-0421.12540

46. Arimitsu H, Inoue K, Sakaguchi Y, Lee J, Fujinaga Y, Watanabe T, Ohyama T, Hirst R, Oguma K. 2003. Purification of fully activated Clostridium botulinum serotype B toxin for treatment of patients with dystonia. Infect Immun 71:1599–1603; PMID:12595486; https://doi.org/10.1128/IAI.71.3.1599-1603.2003

47. Reed L.J., Muench H. 1938. A simple method of estimating fifty per cent endpoints. Am J Hyg 27: 493–497; https://doi.org/10.1093/oxfordjournals.aje.a118408

48. Kondo H, Shimizu T, Kubonoya M, Izumi N, Takahashi M, Sakaguchi G. 1984. Titration of botulinum toxins for lethal toxicity by intravenous injection into mice. Jpn J Med Sci Biol 37: 131–135; PMID:6503025; https://doi.org/10.7883/yoken1952.37.131

